# Preclinical Evaluation of [^18^F]P4B-2412 as Phosphodiesterase 4B Radioligand for Positron Emission Tomography Imaging

**DOI:** 10.1101/2025.01.16.633384

**Authors:** Zhendong Song, Yinlong Li, Siyan Feng, Jiahui Chen, Xin Zhou, Zachary Zhang, Zhenkun Sun, Jian Rong, Chunyu Zhao, Ahmad Chaudhary, Jimmy S. Patel, Yabiao Gao, Thomas L. Collier, Chongzhao Ran, Tyler S. Beyett, Achi Haider, Wito Richter, Hongjie Yuan, Steven H. Liang

## Abstract

Phosphodiesterase 4B (PDE4B) plays a critical role in cAMP hydrolysis and is highly expressed in brain regions associated with neuroinflammation and neurodegenerative disorders. Selective PDE4B radioligands hold significant potential for elucidating disease mechanisms and enabling target occupancy measurements. In this study, we developed [^18^F]P4B-2412, a novel PDE4B-selective radioligand, and evaluated its utility for PET imaging. [^18^F]P4B-2412 was synthesized with high radiochemical yield (27.2%), excellent radiochemical purity (99%), and favorable molar activity (66.2 ± 2.5 GBq/μmol). *In vitro* autoradiography revealed high PDE4B binding, while dynamic PET imaging demonstrated high in vivo specificity for PDE4B in rodent brain regions. Notably, [^18^F]P4B-2412 exhibited excellent metabolic stability *in vitro* and *in vivo*. These findings establish [^18^F]P4B-2412 as a promising PET probe for imaging PDE4B *in vivo*, offering a valuable tool for investigating neuroinflammation and advancing CNS drug development.

## INTRODUCTION

Cyclic nucleotide phosphodiesterases (PDEs) comprise a superfamily of isoenzymes that hydrolyze the second messengers cyclic adenosine monophosphate (cAMP) and cyclic guanosine monophosphate (cGMP).^1-3^ In most mammals, PDEs are encoded by 21 genes, which in turn are grouped into 11 PDE families by sequence homology, with some PDE families encoding for cAMP-specific enzymes (PDE4, PDE7, and PDE8), others for cGMP-specific enzymes (PDE5, PDE6, and PDE9), and some that can hydrolyze both cyclic nucleotides (PDEs 1-3, PDE10, and PDE11). The PDE4 family comprises four genes, PDE4A, PDE4B, PDE4C, and PDE4D, and is arguably the largest and most widely expressed family of cAMP-specific PDEs.^4^

Previous studies have identified PDE4B as the predominant PDE4 isoenzyme in key brain regions, including the thalamus, hypothalamus, cerebellum, and substantia nigra.^5-7^ PDE4B inhibition has been linked to anti-neuroinflammatory and precognitive effects – with potential for the treatment of schizophrenia.^8-10^ Further, Nerandomilast (BI 1015550), a PDE4B-preferrring inhibitor currently investigated in phase III clinical trials, has shown significant efficacy in improving lung function in patients with idiopathic pulmonary fibrosis (IPF).^11^ Several pan-PDE4 inhibitors, such as roflumilast, apremilast, and crisaborole, have been approved for the treatment of chronic obstructive pulmonary disease (COPD), psoriasis, and atopic dermatitis, respectively.^12^ However, the clinical utility of pan- PDE4 inhibitors is often limited by gastrointestinal side effects, including nausea and emesis. These adverse effects may be primarily due to inhibition of PDE4D, given that the genetic deletion of PDE4D, but not the deletion of PDE4B, shortens the duration of alpha_2_-adrenoceptor-mediated anesthesia, a behavioral correlate of emesis, in mice.^13^ Conversely, studies investigating PAN-PDE4 inhibitor– induced gastroparesis and hypothermia in mice as surrogate markers of nausea and emesis suggest that these adverse effects arise from the concurrent inactivation of multiple PDE4 subtypes. In contrast, selective targeting of individual PDE4 subtypes does not appear to elicit these adverse effects.^14, 15^ In either case, the development of selective PDE4B inhibitors represents a promising therapeutic strategy for treating neuroinflammation and CNS diseases, offering an expanded therapeutic window with reduced adverse effects.

Positron emission tomography (PET) imaging plays a pivotal role in preclinical and clinical drug development, enabling disease diagnosis and staging, therapy monitoring as well as target occupancy measurements. Developing highly selective PDE4B radioligands is essential for mapping PDE4B expression and tissue distribution in vivo. (*R*)-[^11^C]rolipram, a pan-PDE4 radioligand, lacks the specificity needed for PDE4B imaging due to low selectivity across other PDE4 isoforms (Figure 1A).^16^ Recently, Pfizer developed [^18^F]PF-06445974, a potent and PDE4B-preferring radioligand, which exhibited reasonable brain permeability and specific binding in non-human primate (NHP) brains (Figure 1B).^17^ In humans, [^18^F]PF-06445974 exhibited high brain uptake (peak SUV: 2-3), with the thalamus showing the highest distribution, consistent with PDE4B expression patterns. However, its slow kinetics necessitates prolonged scanning times, which is suboptimal for clinical applications. Additionally, brain efflux mediated by permeability glycoprotein (P-gp) and breast cancer resistance protein (BCRP) may complicate quantification, warranting further investigation.^18^ The latest study by Zhao *et al*. reported on the development of an effective PET radioligand, [^11^C]ZTP-1, for imaging PDE4B in rat and monkey brains. Its short half-life and high molar activity make it suitable for flexible clinical study design and support its potential as a shorter-lived alternative to [^18^F]PF-06445974 for human PDE4B imaging (Figure 1B).^19^

**Figure 1.**
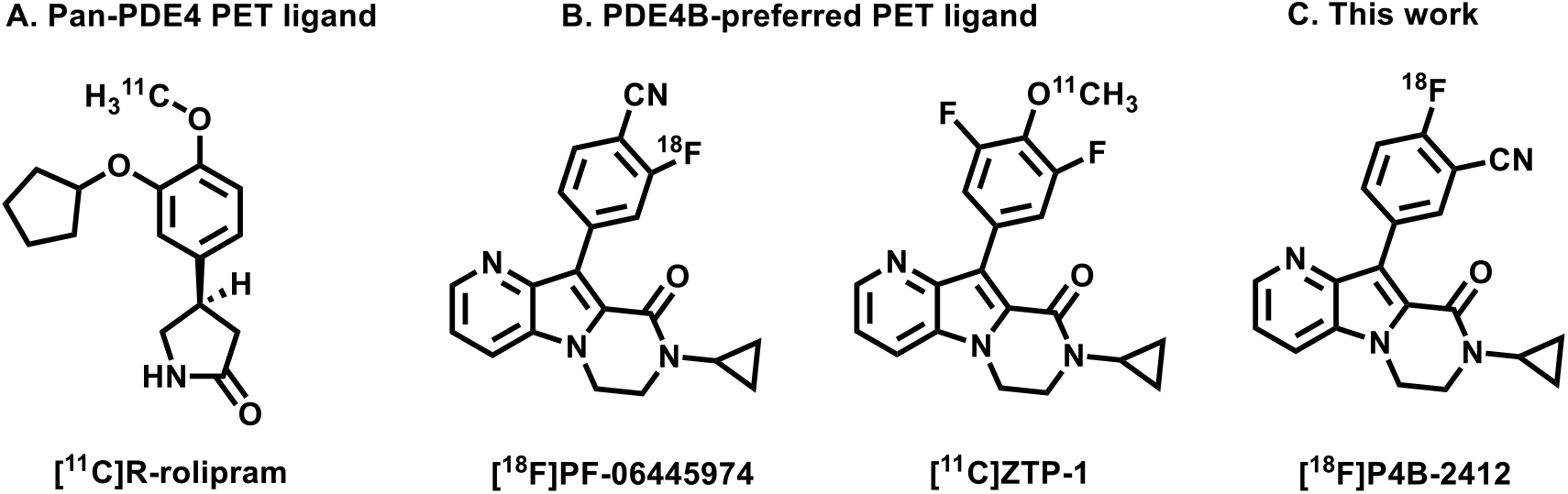
(A) Chemical structure of previously reported pan-PDE4 ligand [^11^C]*R*-rolipram; (B) Chemical structures of PDE4B-targeted PET ligand, [^18^F]PF-06445974 and [^11^C]ZTP-1; (C) Chemical structure of [^18^F]P4B-2412 (this work).

As part of the efforts to develop next-generation PDE4B-selective radioligands for neuro-PET imaging, a series of selective PDE4B inhibitors, based on tricyclic pyrrolopyridine scaffold, were recently reported, as exemplified by PF-06445974 and compound P4B-2412.^17^ As a regioisomer of PF- 06445974, compound P4B-2412 incorporates a *p*-fluoro *m*-cyano phenyl group and exhibits high PDE4B inhibitory potency (IC_50_ = 2 nM) with 22-fold selectivity over PDE4D (Figure 1C). Although its *in vitro* selectivity is moderate, we hypothesized that improved washout kinetics may be achieved without compromising PDE4B/PDE4D selectivity *in vivo*. Consequently, in this study, we synthesized [^18^F]P4B-2412 and characterized its performance characteristics by in vitro autoradiography, PET imaging, whole-body *ex vivo* biodistribution, and stability assessments *in vivo* and *in vitro*.

## RESULTS AND DISCUSSION

### Chemical Synthesis and Pharmacology

The reference compound P4B-2412 was synthesized according to a previously reported methods, with minor modifications.^17^ Specifically, commercially available intermediate **1** was reacted with **2** *via* a Suzuki coupling reaction, yielding the final product P4B-2412 in 66% yield, as illustrated in Scheme 1.

**Scheme 1.**
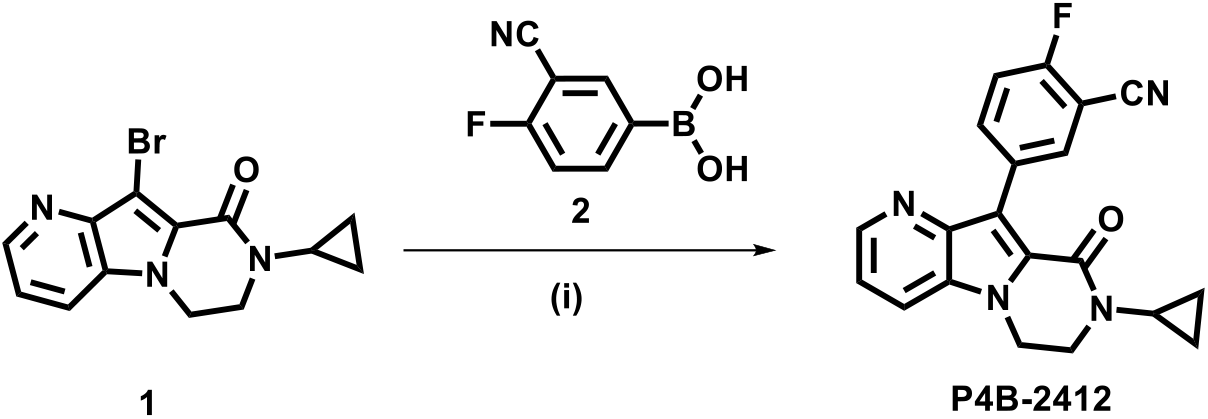
Synthesis of reference compound P4B-2412^a^ ^a^Reagents and conditions: (i) Pd(PPh_3_)_4_, K_2_CO_3_, dioxane/H_2_O (1 mL / 0.3 mL), N_2_, microwave 130 ^o^C, 1 h, 66% yield.

P4B-2412 exhibited favorable properties for CNS-targeting, characterized by a high MPO score (4.0)^20^, a low logBB value (-0.415), and positive CNS activity predictions using QikProp.^21^ The topological polar surface area (tPSA) was calculated by ChemDraw and the logD value (2.38) was experimentally determined using the “shake flask method”.^22^ Of note, P4B-2412 exhibited high selectivity over other PDE families including PDE1A, PDE1B, PDE1C, PDE2A, PDE3A, PDE3B, PDE5A, PDE8A, PDE9A, and PDE10A with the inhibitory activities exceeding 3 μM. Collectively, these findings suggest that P4B-2412 possesses a high binding affinity to PDE4B and favorable physicochemical properties, aligning with CNS PET design parameters (Figure 2).

**Figure 2.**
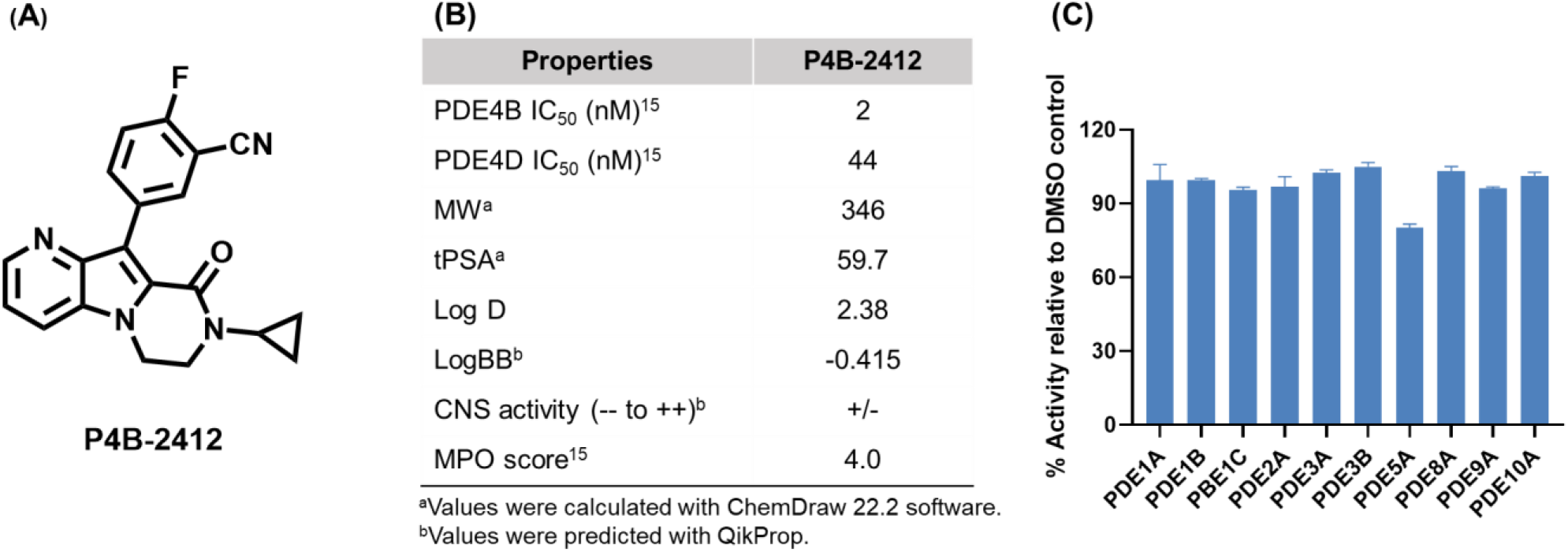
The structure and pharmacological profiles of compound P4B-2412. (A) Structure of P4B-2412. (B) Representative pharmacological and physicochemical properties of P4B-2412. (C) Inhibition of other PDEs by compound P4B-2412 at 3 µM. All data are mean ± SD, n ≥ 2.

### Radiochemistry

To leverage the activating effect of the *m*-cyano group on the phenyl ring, we synthesized precursor **5** with a *p*-nitro substituent. As outlined in Scheme 2, intermediate **4** was prepared by a Suzuki coupling reaction between commercially available **3** and bis(pinacolato)diboron, followed by a second Suzuki coupling reaction with intermediate **1** to give the nitro precursor **5**. ^18^F-labeling of **5** proceeded *via* a nucleophilic substitution reaction using [^18^F]fluoride/Kryptofix222 in DMF at 130 ^o^C for 10 min, as previously reported method.^17, 23-25^ [^18^F]P4B-2412 was obtained with a good radiochemical yield (non-decay-corrected RCY: 27.2%) and an excellent molar activity of 66.2 ± 2.5 GBq/μmol. The radiochemical purity of [^18^F]P4B-2412 exceeded 99%, rendering it suitable for further evaluation.

**Scheme 2.**
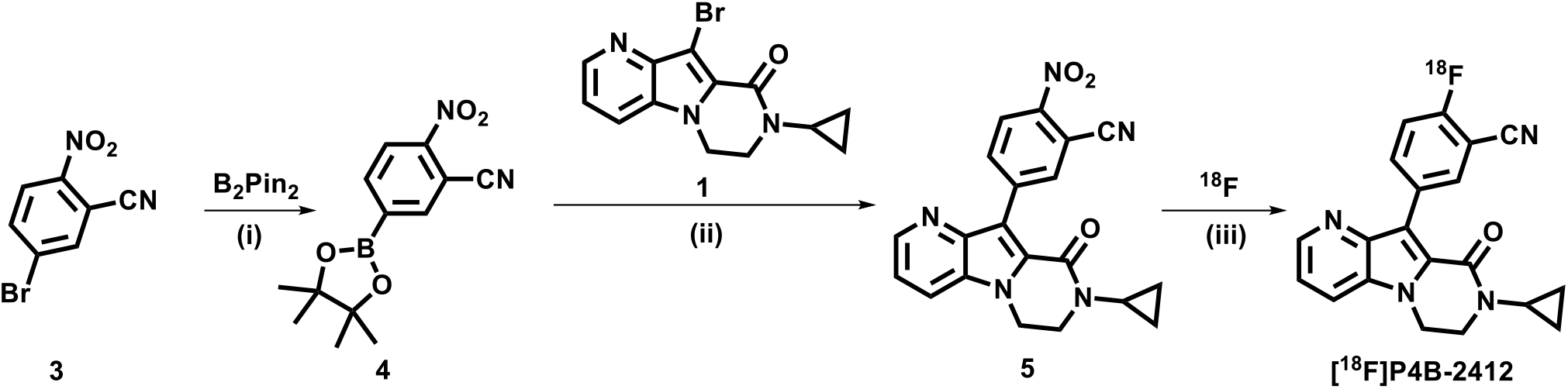
Synthesis of nitro precursor **5** and radiosynthesis of ligand [^18^F]P4B-2412^a^ ^a^Reagents and conditions: (i) B_2_Pin_2_, Pd(dppf)Cl_2_, KOAc, dioxane, N_2_, 100 ^o^C, 3 h. (ii) Pd(dppf)Cl_2_, K_2_CO_3_, dioxane/H_2_O (3/1, v/v), N_2_, 100 ^o^C, 2 h, 47% yields of two steps. (iii) [^18^F]fluoride/Kryptofix 222, K_2_CO_3_, DMF, 130 °C, 10 min, 27.7% non-decay-corrected RCY, 66.2 ± 2.5 GBq/μmol molar activity.

### *In Vitro* Autoradiography

To evaluate the specific binding of [^18^F]P4B-2412 to PDE4B, an *in vitro* autoradiography study was conducted using rat brain sections. [^18^F]P4B-2412 demonstrated excellent brain uptake across various brain regions, with the highest radioactivity accumulation observed in the thalamus, followed by the cortex, hippocampus, striatum, and cerebellum (Figure 3). Co-incubation with rolipram (a PDE4B/4D inhibitor, 10 μM) and PF-06445974 (a selective PDE4B inhibitor, 10 μM) diminished [^18^F]P4B-2412 binding in all brain regions. In contrast, treatment of a high dose of selective PDE4D inhibitor zatolmilast (10 μM) had minimal effect on [^18^F]P4B-2412 binding, suggesting that [^18^F]P4B-2412 preferentially targeted PDE4B rather than PDE4D across the rat brain.

**Figure 3.**
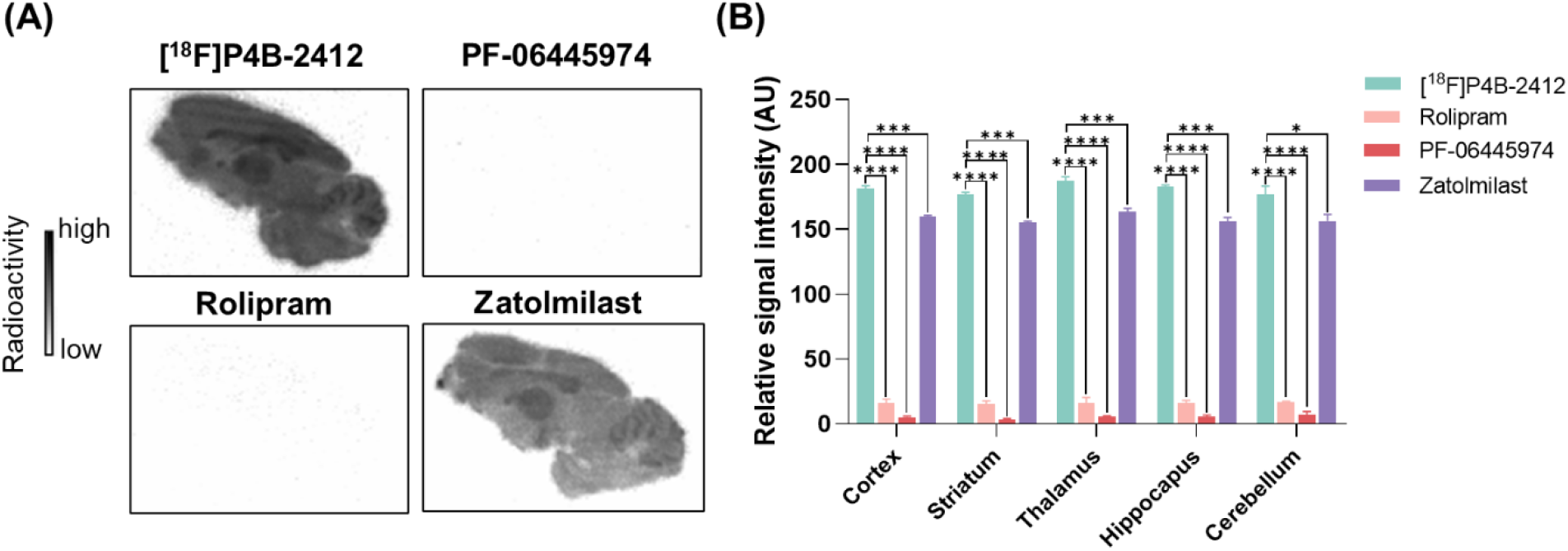
*In vitro* autoradiography studies with [^18^F]P4B-2412 in rat brain sections. (A) Representative images in baseline and blocking (rolipram, PF-06445974, and zatolmilast, 10 μM) studies; (B) Quantification of *in vitro* autoradiography studies with [^18^F]P4B-2412. All data are reported as mean ± SD, n = 3. \**P* ≤0.05, \*\*\**P* ≤ 0.001, \*\*\*\**P* ≤ 0.0001.

### PET Imaging Studies in Mice Brains

To evaluate whether [^18^F]P4B-2412 exhibits specific binding to PDE4B *in vivo*, dynamic PET imaging studies were performed in CD1 mice. At baseline, [^18^F]P4B-2412 demonstrated reasonable brain uptake, with high accumulation observed in the cortex, striatum, thalamus, and hippocampus, consistent with the distribution of PDE4B in the mouse brain (Figure 4A, 4B). The baseline time–activity curves (TACs) revealed that [^18^F]P4B-2412 reached peak uptake at five minutes (standardized uptake value, SUV ∼ 0.45), followed by a gradual washout over 60 minutes across all brain regions. To further assess the reversibility of [^18^F]P4B-2412 binding *in vivo*, we next carried out a “chase” experiment using PF-06445974 (3 mg/kg) administered 15 minutes post-ligand injection. The brain uptake significantly decreased after the injection of PF-06445974, confirming a reversible binding characteristic of [^18^F]P4B-2412 toward PDE4B (Figure 4D). In blocking studies, pretreatment with rolipram and PF-06445974, administered 10 min prior to radioligand injection significantly reduced radioactivity levels in the whole brain. In contrast, the selective PDE4D inhibitor zatolmilast (3 mg/kg) showed no blocking effect (Figure 4C). Although P4B-2412 has been shown to exhibit moderate selectivity for PDE4B over PDE4D *in vitro* (22-fold), PET imaging demonstrated that [^18^F]P4B-2412 possesses significantly higher selectivity for these two subtypes *in vivo*. [^18^F]P4B-2412 achieves more selective binding to PDE4B over PDE4D, despite its only moderate *in vitro* selectivity.

**Figure 4.**
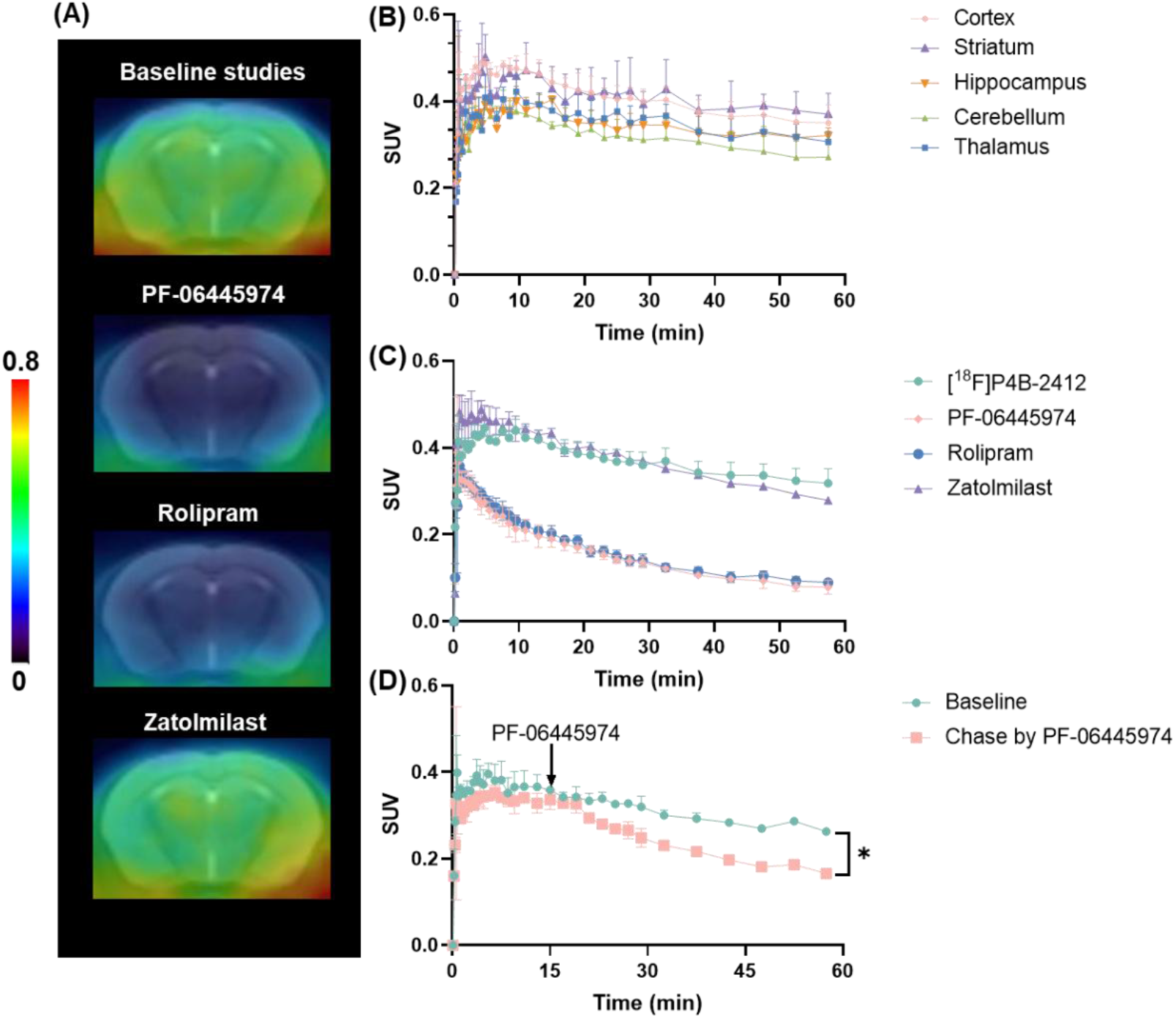
PET imaging studies of [^18^F]P4B-2412 in mice brains. (A) Representative summed (0−60 min) PET images with [^18^F]P4B-2412 under baseline and blocking (rolipram: 3 mg/kg, PF-06445974: 3 mg/kg, zatolmilast: 3 mg/kg) conditions, all blockers were injected *via* tail vein of mice 10 minutes prior to tracer injection. (B) TACs of [^18^F]P4B-2412 in brain regions. (C) TACs of [^18^F]P4B-2412 in the whole-brain under baseline and blocking studies. (D) Chase studies of [^18^F]P4B- 2412 with PF-06445974 (3 mg/kg) at 15 min post tracer administration in mice. All data is presented as mean ± SD, n ≥ 3.

### *Ex Vivo* Biodistribution Studies in CD1 Mice

To further investigate the *in vivo* distribution properties of [^18^F]P4B-2412, *ex vivo* whole-body biodistribution studies were performed in CD1 mice. As shown in Figure 5, radioactivity levels in various organs were measured at 5, 15, 30, and 60 minutes post-radioligand administration. At 5 minutes, high radioactivity accumulation was observed in the heart, pancreas, small intestine, kidneys, and liver, with moderate uptake in the brain. At later time points, radioactivity levels remained high in the small intestine, liver, and kidneys, indicating hepatobiliary and urinary elimination pathways. Notably, the low uptake of [^18^F]P4B-2412 in the bone suggests minimal radiodefluorination, supporting its metabolic stability *in vivo*.

**Figure 5.**
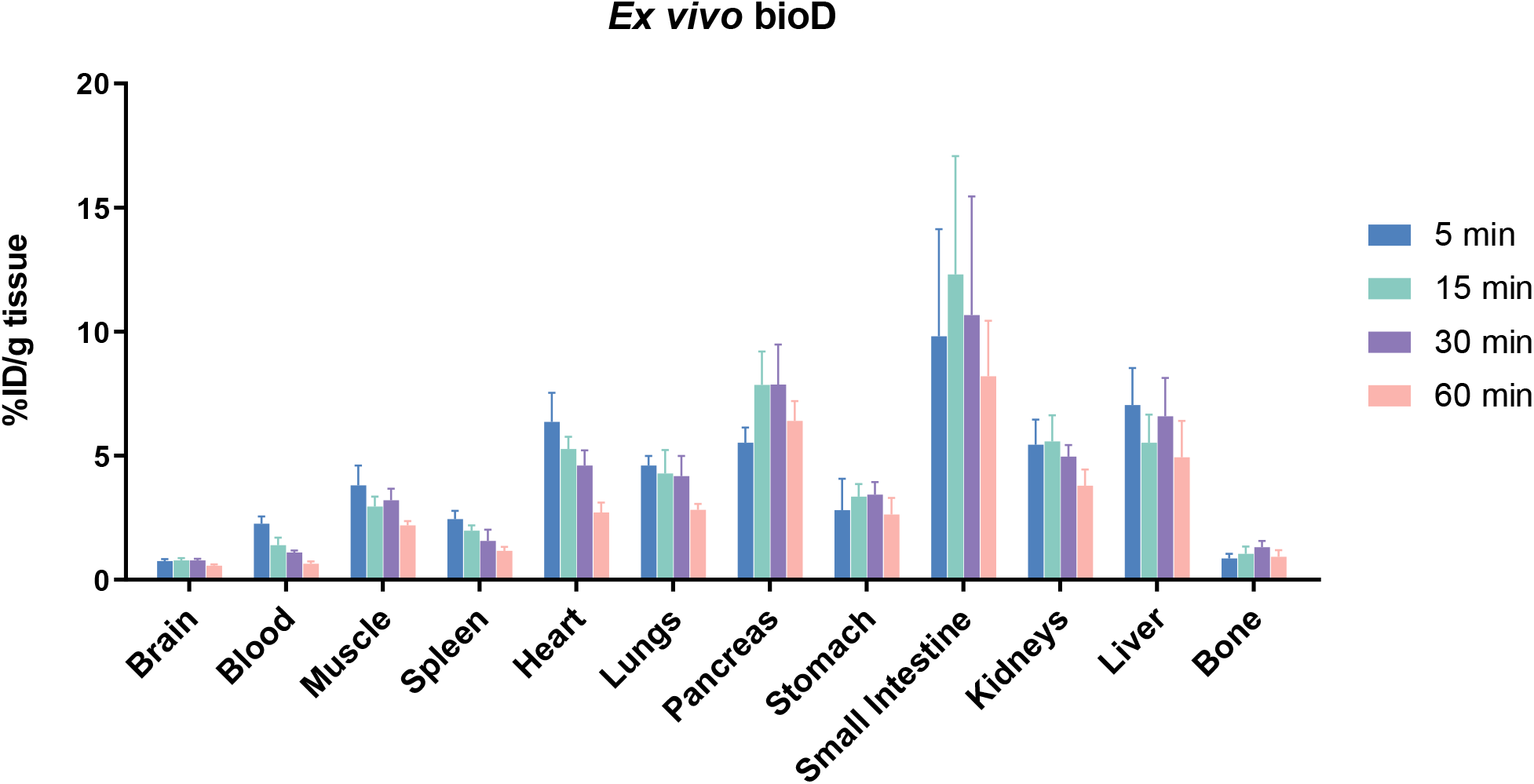
*Ex vivo* biodistribution studies in CD1 mice. All data is provided as mean ± SD, n = 4.

### Stability of [^18^F]P4B-2412 *in Vitro* and *in Vivo*

The *in vitro* stability of [^18^F]P4B-2412 was evaluated in the serum and microsomes from mice, rats, NHPs, and humans (Figure 6). [^18^F]P4B-2412 exhibited excellent stability in serum across all species, with over 98% of parent radioligands remining intact (Figure 6A). Furthermore, [^18^F]P4B-2412 demonstrated robust stability in microsomal assays during a 90-min incubation (Figure 6B). Subsequently, *ex vivo* metabolic analysis of [^18^F]P4B-2412 was performed with mouse brain and plasma samples (Figure 7). At 30 minutes post-administration, 95.4% of the parent radioligand was detected in the brain, while 28.6% remained intact in the plasma. These findings suggest [^18^F]P4B- 2412 possesses reasonable *in vivo* stability.

**Figure 6.**
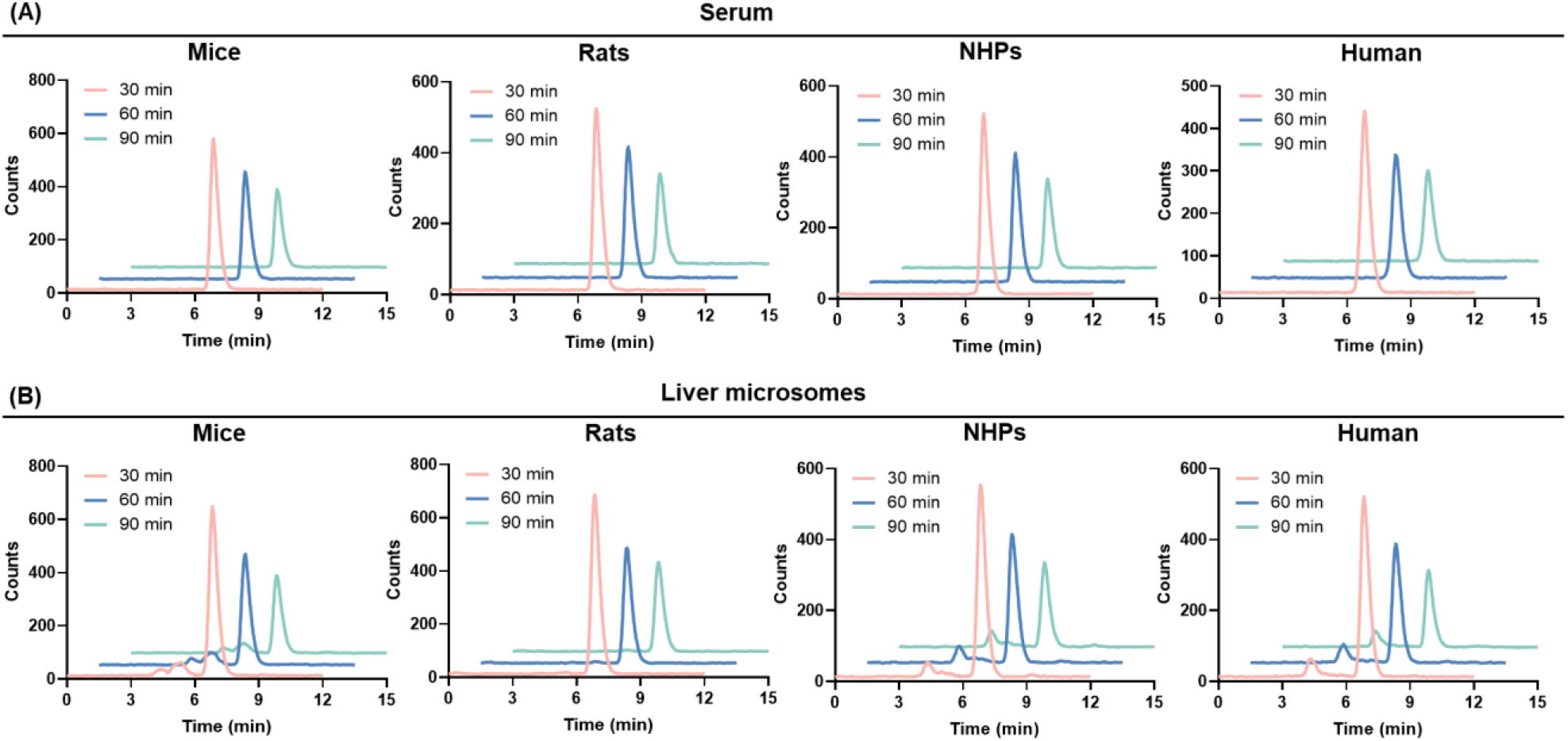
(A) Stability of [^18^F]P4B-2412 in serum of mice, rats, NHPs, and humans at 30, 60, and 90 min; (B) Stability of [^18^F]P4B-2412 in liver microsomes from mice, rats, NHPs, and humans at 30, 60, and 90 min.

**Figure 7.**
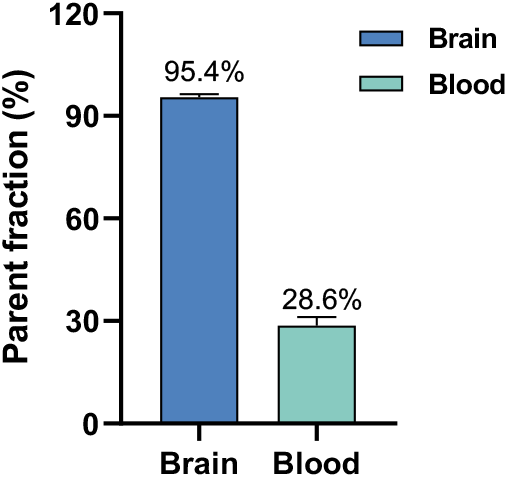
Radiometabolic stability in the brain and plasma of CD1 mice 30-minute post injection of [^18^F]P4B-2412. All data are presented as mean ± SD, n = 2.

## CONCLUSIONS

PDE4B inhibition has emerged as a promising therapeutic strategy with implications in neurodegenerative disorders. As such, the development of a selective PDE4B PET radioligand remains an unmet clinical need. We herein developed a fluoride-18 labeling PDE4B PET radioligand, [^18^F]P4B- 2412. [^18^F]P4B-2412 was obtained in high radiochemical yield and favorable molar activities, as well as radiochemical purities. *In vitro* autoradiography demonstrated distribution patterns in accordance with known PDE4B expression patterns. PET imaging in mice brain revealed significantly enhanced specificity, with negligible binding with PDE4D as confirmed by blocking with a selective PDE4D inhibitor, zatolmilast. Collectively, these finding highlights an improved *in vivo* selectivity and underscore the potential of [^18^F]P4B-2412 as a PDE4B imaging agent. Furthermore, [^18^F]P4B-2412 showed rapid brain uptake in the mice, followed by a gradual washout, suggesting future efforts should focus on improving the washout kinetics to further enhance signal clarity and minimize background noise. [^18^F]P4B-2412 demonstrated excellent metabolic stability in the brain, however, the brain uptake was hampered by P-gp and/or BCRP efflux liabilities. These findings highlight the importance of minimizing transporter-mediated clearance in future tracer development efforts. In summary, [^18^F]P4B-2412 represents a significant advancement in the development of PDE4B-selective PET tracers. Improving efflux liabilities and demonstrating suitable performance characteristics in NHPs will be critical steps toward clinical implementation.

## MATERIALS AND METHODS

### Chemical Synthesis

#### General Information

The experimental procedures were conducted as previous reports with minor modifications. All reagents and solvents were purchased from commercial sources without further purification. ^1^H NMR spectra were recorded on a 400 MHz Bruker spectrometer. Multiplicity of ^1^H NMR signals was reported as singlet (s), doublet (d), triplet (t), quartert (q), and multiplet (m), br (broad signal), dd (doublet of doublets), and so forth. Coupling constants (J) are reported in hertz (Hz). Mass spectra (MS) was recorded on a Shimadzu LC/MS-2020 spectrometer. The preparation of radioligands was performed by reverse-phase HPLC on Phenomenex C18 column (10.0 × 250 mm, 5 μM). The purity of radioligands was confirmed by Agilent 1100 series HPLC system on Waters XSELECT HSS T3 column (4.6 × 150 mm, 5 μM). All animal studies were carried out in accordance with ethical guidelines of Institutional Animal Care and Use Committee (IACUC) of Emory University (PROTO202200003). CD-1 mice (female, 22−24 g, 5−6 weeks) were fed *ad libitum* with food and water under a condition of 12 h light/12 h dark cycle.

### Synthesis of PDE4B Inhibitor P4B-2412

#### 5-(8-cyclopropyl-9-oxo-6,7,8,9-tetrahydropyrido[2’,3’:4,5]pyrrolo[1,2-a]pyrazin-10-yl)-2- fluorobenzonitrile (P4B-2412)

To a solution of 10-bromo-8-cyclopropyl-7,8-dihydropyrido[2’,3’:4,5]pyrrolo[1,2-a]pyrazin-9(6H)- one **1** (60 mg, 0.24 mmol) and (3-cyano-4-fluorophenyl)boronic acid **2** (42.8 mg, 0.26 mmol) in dioxane/H_2_O (1 mL/0.3 mL) were added K_2_CO_3_ (99.4 mg, 0.72 mmol) and Pd(PPh_3_)_4_ (13.9 mg, 0.012 mmol) under N_2_. The resulting mixture was heated to 130 ^o^C for 1 h by Microwave. The reaction was concentrated and purified by flash chromatography (DCM/EA=10/1) to give P4B-2412 (45 mg, 66% yield) as a yellow solid. LC/MS: 347.2 [M+H]^+^, HPLC purity: 97%. ^1^H NMR (400 MHz, CDCl_3_) *δ* 8.61 (d, *J* = 4.4 Hz, 1H), 8.12 (dd, *J* = 6.3, 2.2 Hz, 1H), 8.11 – 8.05 (m, 1H), 7.76 – 7.68 (m, 1H), 7.34 (dd, *J* = 8.4, 4.5 Hz, 1H), 7.29 (d, *J* = 8.8 Hz, 1H), 4.32 (dd, *J* = 6.9, 4.6 Hz, 2H), 3.91 (dd, *J* = 6.8, 4.6 Hz, 2H), 2.85 (tt, *J* = 7.2, 4.0 Hz, 1H), 0.98 (td, *J* = 7.2, 5.5 Hz, 2H), 0.81 – 0.71 (m, 2H).

### Synthesis of Precursor 5

#### 2-nitro-5-(4,4,5,5-tetramethyl-1,3,2-dioxaborolan-2-yl)benzonitrile (4)

To a solution of 5-bromo-2-nitrobenzonitrile (1.00 g, 4.40 mmol), bis (pinacolato)diborane (2.24 g, 8.81 mmol), and potassium acetate (1.30 g, 13.21 mmol) in dioxane (10 mL) was added Pd(dppf)Cl_2_ (100 mg, 0.44 mmol). The mixture was heated at 100 ^o^C for 3 h. The mixture was concentrated, and the crude product was passed through a plug of silica gel, eluting with DCM (20 mL*4). The eluent was then concentrated to give 2-nitro-5-(4,4,5,5-tetramethyl-1,3,2-dioxaborolan-2-yl)benzonitrile **4** (1 g, crude) without further purification.

#### 5-(8-cyclopropyl-9-oxo-6,7,8,9-tetrahydropyrido[2’,3’:4,5]pyrrolo[1,2-a]pyrazin-10-yl)-2- nitrobenzonitrile (5)

To a solution of 10-bromo-8-cyclopropyl-7,8-dihydropyrido[2’,3’:4,5]pyrrolo[1,2-a]pyrazin-9(6*H*)- one **1** (150 mg, 0.59 mmol) and 2-nitro-5-(4,4,5,5-tetramethyl-1,3,2-dioxaborolan-2-yl)benzonitrile **4** (162 mg, 0.49 mmol) in dioxane/H_2_O (3 mL/1 mL) were added K_2_CO_3_ (203 mg, 1.47 mmol) and Pd(dppf)Cl_2_ (39 mg, 0.049 mmol) under N_2_. The mixture was heated to 100 ^o^C for 2 h. The reaction was concentrated and purified by flash chromatography (DCM/EA=10/1) to give compound **5** (93 mg, 47% for 2 steps) as a yellow solid. LC/MS: 374.1[M+H]^+^, HPLC purity: > 99%. ^1^H NMR (400 MHz, CDCl_3_) *δ* 8.64 (d, *J* = 3.4 Hz, 1H), 8.47 (s, 1H), 8.36 (t, *J* = 7.3 Hz, 2H), 7.78 (d, *J* = 7.7 Hz, 1H), 7.39 (s, 1H), 4.36 (s, 2H), 3.95 (s, 2H), 2.88 (dt, *J* = 7.0, 3.4 Hz, 1H), 1.00 (q, *J* = 6.5 Hz, 2H), 0.84 – 0.74 (m, 2H).

### Radiosynthesis of Radioligand [^18^F]P4B-2412

The [^18^F]F^-^ was trapped by a Sep-Pak QMA Plus Light cartridge (Waters) and was eluted into a vial with a solution of K_2_CO_3_ (2 mg) in 0.5 mL of water, followed by a solution of Kryptofix 222 (10 mg) in 1 mL of MeCN. After drying three times using MeCN at 110 ^o^C under a N_2_ atmosphere, the reaction mixture was dissolved in a solution of precursor **5** (1.5 mg) in DMF (0.4 mL) and heated at 130 ^o^C for 10 min. The reaction was cooled to room temperature and mobile phase (4 mL) was added (35% MeCN/0.1 M ammonium acetate). The solution was purified by semi prep-HPLC (Lablogic, Phenomenex Luna C-18 column [250 × 10 mm, 5 μm], mobile phase 35% MeCN/0.1 M ammonium acetate, flow: 5 mL/min) and then collected [^18^F]P4B-2412 in a sterile vial with the retention time of 22 min. The fractions were diluted with water (35 mL) and trapped on a Sep-Pak light C18 cartridge (Waters). The final radiotracer was washed with ethanol (0.3 mL) and formulated with saline for further studies.

### Measurement of Log D

The Log D value was determined using previously established methods.^26^ 2 μCi of [^18^F]P4B-2412 was added to a pre-mixed solution of PBS (pH 7.4, 3 mL) and n-octanol (3 mL), followed by vortexing for 5 minutes. Subsequently, 0.8 mL of both the PBS and n-octanol layers were transferred to a centrifuge tube and centrifuged at 12,000 rpm for 5 minutes. The separated PBS and n-octanol phases were weighed, and their radioactivity was measured using a gamma counter (PerkinElmer Wizard). The Log D value for [^18^F]P4B-2412 was then calculated using the following formula:

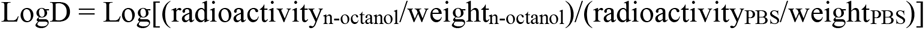

### Autoradiography Studies

Rat brain slides were warmed to room temperature and pre-incubated for 10 minutes in Tris-HCl buffer (50 mM, containing 0.1% bovine serum albumin). The slides were then co-incubated with [^18^F]P4B- 2412 under baseline conditions (with DMSO) or in the presence of rolipram (10 μM) and PF-06445974 (10 μM) for 30 minutes. Following incubation, the slides were rinsed three times in Tris-HCl buffer (50 mM) for 5 minutes each and briefly dipped in water twice (5 seconds per dip). After air-drying under cool conditions, the slides were exposed to a Phosphor Screen (BAS IP MS2025, Cytiva) overnight. Radioactivity was subsequently quantified using an Amersham Typhoon 5 (Cytiva), with data analysis performed in ImageJ software.

### PET imaging Studies in Mice

PET imaging was performed as described in previous studies.^26^ CD1 mice (5–6 weeks old) were anesthetized with 1.5% (v/v) isoflurane throughout the scan. A dose of 80 μCi of [^18^F]P4B-2412 was administered *via* intravenous injection into the tail vein, followed by a 60-minute dynamic scan using a PET scanner (MOLECUBES, beta-cube). For blocking studies, rolipram (3 mg/kg) and PF-06445974 (3 mg/kg) were injected 10 minutes before the PET radioligands. Data reconstruction by PET scanner and analysis using PMOD 4.3 software.

### *Ex Vivo* Whole-body Biodistribution

CD1 mice (5–6 weeks old) received an intravenous injection of 20 μCi of [^18^F]P4B-2412 in 0.1 mL saline *via* the tail vein. At 5-, 15-, 30-, and 60-minutes post-injection, the mice were euthanized, and their organs were harvested and weighed. Radioactivity in each organ was quantified using a gamma counter (PerkinElmer Wizard).

### Stability in Serum and Microsomes

Serum samples from mice, rats, NHPs, and humans were brought to room temperature, and 300 μCi of [^18^F]P4B-2412 was added to 0.4 mL of each. The mixtures were incubated at 37 °C, and at 30, 60, and 90 minutes, 0.1 mL mixture was transferred to 0.2 mL of cold acetonitrile. After vortexing for 15 seconds, the samples were centrifuged at 10,000 g for 5 minutes. The resulting supernatant was analyzed by HPLC using an Agilent 1100 system and a Phenomenex Luna C-18 column (4.6 × 150 mm), with a mobile phase of 25% MeCN/0.1% TFA in water at a flow rate of 1 mL/min. For microsome studies, the reaction mixture contained 0.5 mg/mL microsomes, 300 μCi of [^18^F]P4B-2412, 0.35 mL PBS, 0.05 mL of NADPH regenerating system A, and 0.01 mL of system B. After incubation at 37 °C, the subsequent steps for sample preparation and analysis were identical to those used for serum samples.

### *In Vivo* Metabolite Analysis

CD1 mice (5–6 weeks old) were intravenously administered 300 μCi of [^18^F]P4B-2412 in 0.1 mL saline *via* the tail vein. At 30 minutes post-injection, the mice were euthanized, and their brains and blood were rapidly collected and homogenized with cold acetonitrile. After centrifugation, the supernatants were analyzed using pre-HPLC (Lablogic HPLC) equipped with a Phenomenex Luna C- 18 column (250 × 10 mm, 5 μm). The mobile phase consisted of 45% MeCN/0.1 M ammonium acetate, (flow: 5 mL/min). Fractions were collected every 30 seconds over a 15-minute period, and the radioactivity of each fraction was measured using a gamma counter (PerkinElmer Wizard). Data was analyzed with GraphPad Prism 8.0.

## AUTHOR INFORMATION

### Authors

**Zhendong Song** – *Department of Radiology and Imaging Sciences, Emory University, 1364 Clifton Road, Atlanta, GA 30322, USA*.

**Yinlong Li** – *Department of Radiology and Imaging Sciences, Emory University, 1364 Clifton Road, Atlanta, GA 30322, USA*.

**Siyan Feng** – *Department of Radiology and Imaging Sciences, Emory University, 1364 Clifton Road, Atlanta, GA 30322, USA*

**Jiahui Chen** – *Department of Radiology and Imaging Sciences, Emory University, 1364 Clifton Road, Atlanta, GA 30322, USA*

**Xin Zhou** – *Department of Radiology and Imaging Sciences, Emory University, 1364 Clifton Road, Atlanta, GA 30322, USA*

**Zachary Zhang** – *Department of Radiology and Imaging Sciences, Emory University, 1364 Clifton Road, Atlanta, GA 30322, USA*

**Zhenkun Sun** – *Department of Pharmacology and Chemical Biology, Emory University School of Medicine, Atlanta, GA, 30322, USA*

**Jian Rong** – *Department of Radiology and Imaging Sciences, Emory University, 1364 Clifton Road, Atlanta, GA 30322, USA*

**Chunyu Zhao** – *Department of Radiology and Imaging Sciences, Emory University, 1364 Clifton Road, Atlanta, GA 30322, USA*

**Ahmad F. Chaudhary** – *Department of Radiology and Imaging Sciences, Emory University, 1364 Clifton Road, Atlanta, GA 30322, USA*

**Jimmy S. Patel** – *Department of Radiology and Imaging Sciences, Emory University, 1364 Clifton Road, Atlanta, GA 30322; Department of Radiation Oncology, Winship Cancer Institute of Emory University, Atlanta, GA, 30322, USA*.

**Yabiao Gao** – *Department of Radiology and Imaging Sciences, Emory University, 1364 Clifton Road, Atlanta, GA 30322, USA*

**Thomas L. Collier** – *Department of Radiology and Imaging Sciences, Emory University, 1364 Clifton Road, Atlanta, GA 30322, USA*

**Chongzhao Ran** – *Athinoula A. Martinos Center for Biomedical Imaging, Department of Radiology, Massachusetts General Hospital and Harvard Medical School, Boston, Massachusetts, 02114, USA*. **Tyler S. Beyett** – *Department of Pharmacology and Chemical Biology, Emory University School of Medicine, Atlanta, GA, 30322, USA*.

**Achi Haider** – *Department of Radiology and Imaging Sciences, Emory University, 1364 Clifton Road, Atlanta, GA 30322, USA*

**Wito Richter** – *Department of Biochemistry & Molecular Biology, Center for Lung Biology, University of South Alabama College of Medicine, Mobile, AL, 36688, USA*

**Hongjie Yuan** – *Department of Pharmacology and Chemical Biology, Emory University School of Medicine, Atlanta, GA, 30322, USA*.

### Author Contributions

Z.S. and Y.L. contributed equally. All the authors contributed to this manuscript and have approved its final version. Z.S. and S.H.L. designed the study and prepared the manuscript. Z.S., Y.L., S.F., J.C., X.Z., Z.Z., Z.S., J.R., C.Z., A.F.C., J.S.P., Y.G., T.L.C., C.R., T.S.B., A.H., W.B. and H.Y. performed and analyzed experiments. S.L. conceived the project and revised the manuscript.

### Notes

The authors declare no competing financial interest.

## ACKNOWLEDGMENTS

We thank Emory Center for Systems Imaging Radiopharmacy (Ronald J. Crowe, RPh, BCNP; Karen Dolph, RPh; M. Shane Waldrep) & Department of Radiology and Imaging Sciences, Emory University School of Medicine for general support. We also thank the National Institute of Mental Health’s Psychoactive Drug Screening Program (NIMH PDSP) for compound off-target screening. J.S.P. is supported by NCI T32CA275777. S.H.L. gratefully acknowledges the support provided, in part, by the NIH grants (AG078058, AG079956, P30AG066511, and S10OD034326, United States), Emory Radiology Chair Fund, and Emory School of Medicine Endowed Directorship.

## REFERENCES

(1) Bender, A. T.; Beavo, J. A. Cyclic nucleotide phosphodiesterases: molecular regulation to clinical use. Pharmacol Rev 2006, 58 (3), 488–520.

(2) Azam, M. A.; Tripuraneni, N. S. Selective Phosphodiesterase 4B Inhibitors: A Review. Sci Pharm 2014, 82 (3), 453–481.

(3) Maurice, D. H.; Ke, H.; Ahmad, F.; Wang, Y.; Chung, J.; Manganiello, V. C. Advances in targeting cyclic nucleotide phosphodiesterases. Nat Rev Drug Discov 2014, 13 (4), 290–314.

(4) Song, Z.; Huang, Y. Y.; Hou, K. Q.; Liu, L.; Zhou, F.; Huang, Y.; Wan, G.; Luo, H. B.; Xiong, X. F. Discovery and Structural Optimization of Toddacoumalone Derivatives as Novel PDE4 Inhibitors for the Topical Treatment of Psoriasis. J Med Chem 2022, 65 (5), 4238–4254.

(5) Pérez-Torres, S.; Miró, X.; Palacios, J. M.; Cortés, R.; Puigdoménech, P.; Mengod, G. Phosphodiesterase type 4 isozymes expression in human brain examined by in situ hybridization histochemistry and[3H]rolipram binding autoradiography. Comparison with monkey and rat brain. J Chem Neuroanat 2000, 20 (3-4), 349–374.

(6) Wu, Y.; Li, Z.; Huang, Y. Y.; Wu, D.; Luo, H. B. Novel Phosphodiesterase Inhibitors for Cognitive Improvement in Alzheimer’s Disease. J Med Chem 2018, 61 (13), 5467–5483.

(7) Richter, W.; Menniti, F. S.; Zhang, H. T.; Conti, M. PDE4 as a target for cognition enhancement. Expert Opin Ther Targets 2013, 17 (9), 1011–1027.

(8) Su, Y.; Ding, J.; Yang, F.; He, C.; Xu, Y.; Zhu, X.; Zhou, H.; Li, H. The regulatory role of PDE4B in the progression of inflammatory function study. Front Pharmacol 2022, 13, 982130.

(9) Millar, J. K.; Pickard, B. S.; Mackie, S.; James, R.; Christie, S.; Buchanan, S. R.; Malloy, M. P.; Chubb, J. E.; Huston, E.; Baillie, G. S.; et al. DISC1 and PDE4B are interacting genetic factors in schizophrenia that regulate cAMP signaling. Science 2005, 310 (5751), 1187–1191.

(10) McGirr, A.; Lipina, T. V.; Mun, H. S.; Georgiou, J.; Al-Amri, A. H.; Ng, E.; Zhai, D.; Elliott, C.; Cameron, R. T.; Mullins, J. G.; et al. Specific Inhibition of Phosphodiesterase-4B Results in Anxiolysis and Facilitates Memory Acquisition. Neuropsychopharmacology 2016, 41 (4), 1080–1092.

(11) Richeldi, L.; Azuma, A.; Cottin, V.; Kreuter, M.; Maher, T. M.; Martinez, F. J.; Oldham, J. M.; Valenzuela, C.; Gordat, M.; Liu, Y.; et al. Design of a phase III, double-blind, randomised, placebo-controlled trial of BI 1015550 in patients with idiopathic pulmonary fibrosis (FIBRONEER-IPF). BMJ Open Respir Res 2023, 10 (1).

(12) Spina, D. PDE4 inhibitors: current status. Br J Pharmacol 2008, 155 (3), 308–315.

(13) Robichaud, A.; Stamatiou, P. B.; Jin, S. L.; Lachance, N.; MacDonald, D.; Laliberté, F.; Liu, S.; Huang, Z.; Conti, M.; Chan, C. C. Deletion of phosphodiesterase 4D in mice shortens alpha(2)-adrenoceptor-mediated anesthesia, a behavioral correlate of emesis. J Clin Invest 2002, 110 (7), 1045–1052.

(14) McDonough, W.; Aragon, I. V.; Rich, J.; Murphy, J. M.; Abou Saleh, L.; Boyd, A.; Koloteva, A.; Richter, W. PAN-selective inhibition of cAMP-phosphodiesterase 4 (PDE4) induces gastroparesis in mice. Faseb j 2020, 34 (9), 12533–12548.

(15) Boyd, A.; Aragon, I. V.; Rich, J.; McDonough, W.; Oditt, M.; Irelan, D.; Fiedler, E.; Abou Saleh, L.; Richter, W. Assessment of PDE4 Inhibitor-Induced Hypothermia as a Correlate of Nausea in Mice. Biology (Basel) 2021, 10 (12).

(16) Lourenco, C. M.; Houle, S.; Wilson, A. A.; DaSilva, J. N. Characterization of r-[11C]rolipram for PET imaging of phosphodieterase-4: in vivo binding, metabolism, and dosimetry studies in rats. Nucl Med Biol 2001, 28 (4), 347–358.

(17) Zhang, L.; Chen, L.; Beck, E. M.; Chappie, T. A.; Coelho, R. V.; Doran, S. D.; Fan, K. H.; Helal, C. J.; Humphrey, J. M.; Hughes, Z.; et al. The Discovery of a Novel Phosphodiesterase (PDE) 4B-Preferring Radioligand for Positron Emission Tomography (PET) Imaging. J Med Chem 2017, 60 (20), 8538–8551.

(18) Wakabayashi, Y.; Stenkrona, P.; Arakawa, R.; Yan, X.; Van Buskirk, M. G.; Jenkins, M. D.; Santamaria, J. A. M.; Maresca, K. P.; Takano, A.; Liow, J. S.; et al. First-in-Human Evaluation of (18)F-PF-06445974, a PET Radioligand That Preferentially Labels Phosphodiesterase-4B. J Nucl Med 2022, 63 (12), 1919–1924.

(19) Zhao, Q.; Liow, J. S.; Jee, J. E.; Montero Santamaria, J.; Pamie-George, M.; Morse, C.; Wu, S.; Zoghbi, S. S.; Kim, S. W.; Innis, R. B.; et al. [(11)C]ZTP-1: An Effective Short-Lived Radioligand for PET of Rat and Monkey Brain Phosphodiesterase Type 4 Subtype B. J Nucl Med 2025, 66 (7), 1119–1125.

(20) Wager, T. T.; Hou, X.; Verhoest, P. R.; Villalobos, A. Central Nervous System Multiparameter Optimization Desirability: Application in Drug Discovery. ACS Chem Neurosci 2016, 7 (6), 767–775.

(21) Ioakimidis, L.; Thoukydidis, L.; Mirza, A.; Naeem, S.; Reynisson, J. Benchmarking the reliability of QikProp. Correlation between experimental and predicted values. QSAR & Combinatorial Science 2008, 27 (4), 445–456.

(22) Lombardo, F.; Shalaeva, M. Y.; Tupper, K. A.; Gao, F. ElogD(oct): a tool for lipophilicity determination in drug discovery. 2. Basic and neutral compounds. J Med Chem 2001, 44 (15), 2490–2497.

(23) Deng, X.; Rong, J.; Wang, L.; Vasdev, N.; Zhang, L.; Josephson, L.; Liang, S. H. Chemistry for Positron Emission Tomography: Recent Advances in (11) C-, (18) F-, (13) N-, and (15) O-Labeling Reactions. Angew Chem Int Ed Engl 2019, 58 (9), 2580–2605.

(24) Rong, J.; Haider, A.; Jeppesen, T. E.; Josephson, L.; Liang, S. H. Radiochemistry for positron emission tomography. Nat Commun 2023, 14 (1), 3257.

(25) Krishnan, H. S.; Ma, L.; Vasdev, N.; Liang, S. H. (18) F-Labeling of Sensitive Biomolecules for Positron Emission Tomography. Chemistry 2017, 23 (62), 15553–15577.

(26) Song, Z.; Li, Y.; Dahl, K.; Chaudhary, A.; Sun, Z.; Zhou, X.; Chen, J.; Gao, Y.; Rong, J.; Zhao, C.; et al. Discovery of (18)F Labeled AZD5213 Derivatives as Novel Positron Emission Tomography (PET) Radioligands Targeting Histamine Subtype-3 Receptor. Chembiochem 2024, e202400655.

